# The mevalonate pathway couples lipid metabolism to amino acid synthesis via ubiquinone-dependent redox control

**DOI:** 10.1101/2025.08.10.669191

**Authors:** Thi Thinh Nguyen, Mohit Bansal, Jane Ding, Sunil Sudarshan, Han-Fei Ding

## Abstract

The mevalonate pathway produces sterols and isoprenoids that support cancer cell growth, yet its broader metabolic functions remain incompletely defined. Here, we show that this pathway sustains amino acid biosynthesis by promoting mitochondrial NAD⁺ regeneration through ubiquinone-dependent electron transport. Statin-mediated inhibition of the mevalonate pathway impairs oxidative phosphorylation, lowers the NAD⁺/NADH ratio, and suppresses de novo serine and aspartate synthesis, thereby activating the GCN2–eIF2α–ATF4 amino acid deprivation response. The resulting depletion of serine-derived glycine and one-carbon units, together with reduced aspartate availability, limits purine and pyrimidine nucleotide production. Expression of the bacterial NADH oxidase LbNOX or the alternative oxidase AOX restores NAD⁺ levels and rescues statin-induced growth inhibition. These findings suggest that impaired NAD⁺ regeneration is a key mechanism contributing to the anti-proliferative activity of statins, linking the mevalonate pathway to mitochondrial electron transport– dependent control of amino acid metabolism.

**Significance:** This study identifies the mevalonate pathway as a regulator of amino acid biosynthesis through mitochondrial electron transport–dependent NAD⁺ regeneration and reveals redox disruption as a key mechanism contributing to the anti-proliferative effects of statins.

## Introduction

The mevalonate pathway is a highly conserved metabolic pathway that converts acetyl-CoA into cholesterol and a variety of isoprenoids essential for cell function (Figure 1A) ^1,2^. Cholesterol, which constitutes approximately 25% of plasma membrane lipids ^3^, is crucial for membrane integrity and cell proliferation. Isoprenoid intermediates such as farnesyl pyrophosphate (FPP) and geranylgeranyl pyrophosphate (GGPP) are required for the prenylation and activation of RAS-family GTPases, thereby linking the pathway to oncogenic signaling. Additional isoprenoid derivatives, including dolichol, heme A, and ubiquinone (coenzyme Q), contribute to protein glycosylation, mitochondrial respiration, and redox balance ^4–6^. Owing to its broad impact on lipid homeostasis, protein function, and mitochondrial metabolism, the mevalonate pathway is indispensable for cell growth and survival. However, the full extent of the metabolic activities supported by the mevalonate pathway remains incompletely understood.

**Figure 1.**
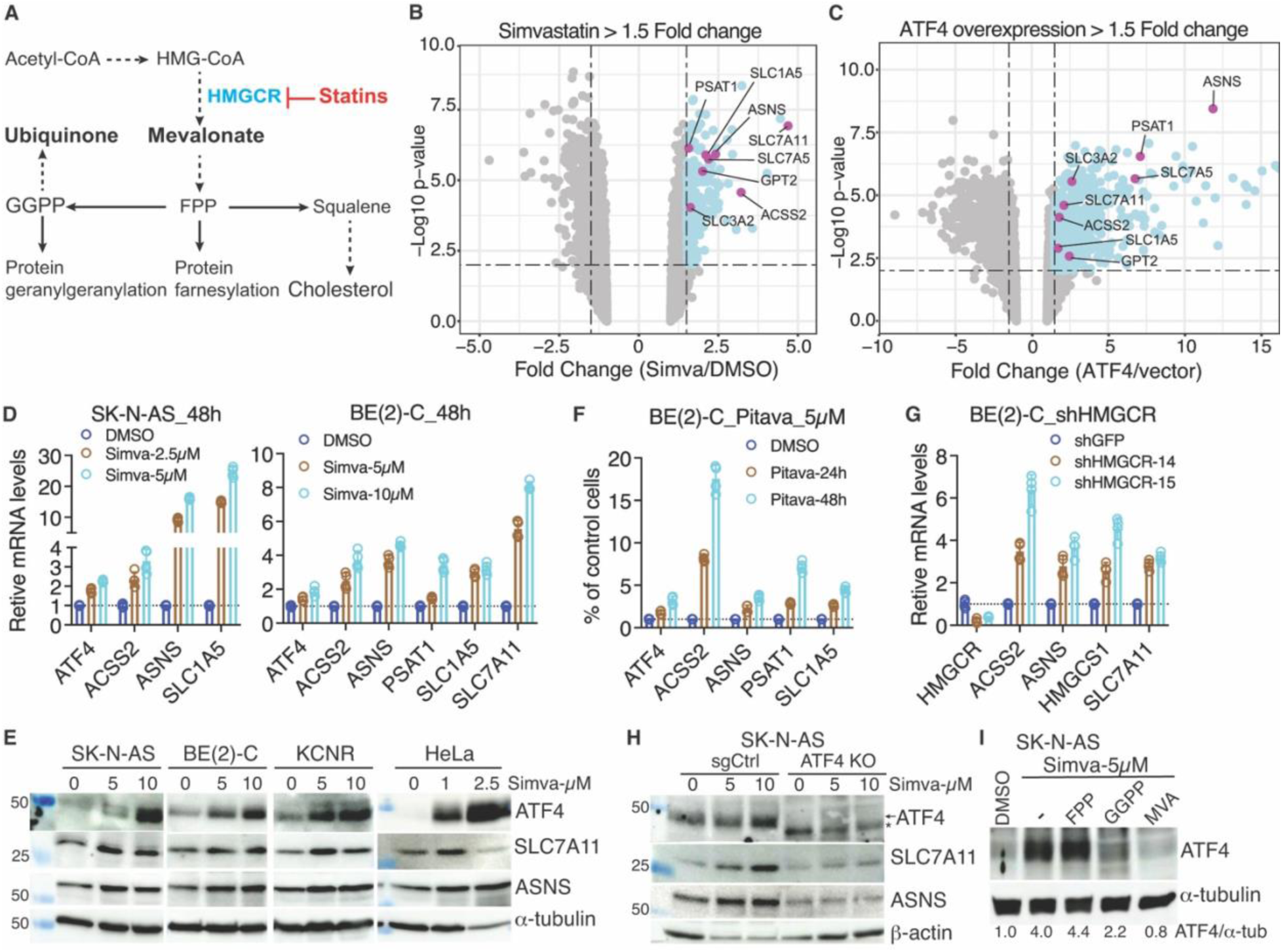
Statins induce an ATF4-dependent adaptive response. (A) Schematic of the mevalonate pathway. (B-C) Volcano plots of microarray gene expression data showing simvastatin (B) and ATF4 overexpression (C) induced a common set of genes involved in amino acid biosynthesis and transport. (D-E) RT-PCR (D) and immunoblotting (E) showing simvastatin induced ATF4, ACSS2, amino acid synthesis enzymes (ASNS, PSAT1) and transporters (SLC1A5, SLC7A11). (F-G) RT-PCR showing pitavastatin (F) and HMRCR knockdown (G) induced the same set of genes. (H) Immunoblotting showing ATF4 knockout (KO) abrogated simvastatin induction of ASNS and SLC7A11. (*) The asterisk indicates a likely truncated ATF4 protein resulting from CRISPR-mediated gene editing. (I) Immunoblotting showing simvastatin-induced ATF4 expression in the absence or presence of supplemental FPP, GGPP, or mevalonate (MVA). ATF4 levels were quantified relative to α-tubulin.

Increased mevalonate metabolism is a common feature of cancer ^2,7^ and has emerged as a potential therapeutic vulnerability ^8–10^. Statins, widely prescribed inhibitors of HMG-CoA reductase (HMGCR) ^11^, the rate-limiting enzyme of the pathway (Figure 1), have demonstrated anti-proliferative effects in multiple cancer models ^8–10^. We previously showed that transcriptional upregulation of the mevalonate pathway is a metabolic feature of high-risk human neuroblastoma and correlates with poor clinical outcome ^12^. More recently, we demonstrated that this upregulation sustains mitotic gene expression and tumor growth in both cell line and patient-derived xenograft models, and that blocking this pathway with statins reduces tumor burden in the TH-MYCN transgenic mouse model of high-risk neuroblastoma ^13^. These findings underscore the critical role of mevalonate metabolism in neuroblastoma development.

In this study, we show that the mevalonate pathway supports amino acid biosynthesis by promoting mitochondrial NAD⁺ regeneration via ubiquinone-dependent electron transport. Inhibition of this pathway impairs oxidative phosphorylation, lowers the NAD⁺/NADH ratio, and selectively suppresses serine and aspartate biosynthesis, triggering the GCN2–eIF2α–ATF4 amino acid deprivation response. Our results establish mitochondrial NAD⁺ regeneration as a key mechanism linking mevalonate metabolism to amino acid biosynthesis and contributing to the growth-suppressive effects of statins.

## Results

### Blocking the mevalonate pathway induces an ATF4-dependent adaptive response

We previously reported that transcriptomic profiling of neuroblastoma SMS-KCNR cells treated with vehicle (DMSO) or simvastatin (5 µM) identified 275 genes upregulated by simvastatin (Table S1) ^13^. These included genes encoding amino acid biosynthetic enzymes (ASNS, GPT2, PSAT1) and amino acid transporters (SLC1A5, SLC3A2, SLC7A5, SLC7A11) (Figure 1B), all of which are established transcriptional targets of ATF4 ^14–16^, a master regulator of amino acid metabolism and stress responses ^17–20^. Consistently, ATF4 overexpression induced transcription of the same set of genes (Figure 1C and Table S2).

We validated these findings by quantitative RT-PCR in two additional neuroblastoma cell lines, SK-N-AS and BE(2)-C (Figure 1D). Immunoblotting confirmed that simvastatin treatment increased protein expression of ATF4 and its targets ASNS, SLC1A5, SLC7A5, and SLC7A11 not only in multiple neuroblastoma cell lines but also in cell lines of different tissue origins, including HeLa (cervical cancer), HCT116 (colon cancer), and 293T (human embryonic kidney) (Figures 1E and S1A). Similar results were obtained with another statin, pitavastatin (Figures 1F and S1B), or by knockdown of HMGCR (Figure 1G), demonstrating that induction of these amino acid metabolic genes is a specific consequence of HMGCR inhibition. Importantly, ATF4 knockout abolished simvastatin-induced expression of these proteins (Figures 1H and S1C).

Supplementation with mevalonate (MVA) fully, and with GGPP partially, reversed simvastatin-induced ATF4 expression (Figure 1I) and rescued the cell growth inhibition by simvastatin (Figure S1D), confirming that these effects arise from blockade of the mevalonate pathway rather than a nonspecific effect of statins.

Collectively, these results demonstrate that inhibition of the mevalonate pathway triggers an ATF4-dependent stress response that is conserved across diverse cell types, indicating a general adaptive mechanism rather than a cell type–restricted phenomenon.

### Blocking the mevalonate pathway activates the amino acid deprivation response

ATF4 is induced by diverse cellular stresses, including amino acid or heme deprivation, endoplasmic reticulum (ER) stress from unfolded proteins, and viral infection ^20,21^. These signals converge on phosphorylation of the α subunit of translation initiation factor 2 (eIF2α), mediated by one of four stress-responsive kinases: HRI (heme deficiency), PKR (viral infection), GCN2 (amino acid deprivation), and PERK (ER stress). Phosphorylation of eIF2α suppresses global protein synthesis while selectively enhancing translation of specific mRNAs, most notably ATF4. Once induced, ATF4 drives a transcriptional program to promote cellular adaptation and survival ^20–22^.

Gene set enrichment analysis (GSEA) of our transcriptomic data revealed that simvastatin treatment activated the amino acid deprivation response (AAR) signature (Figures 2A and 2B). This effect was reversed by mevalonate supplementation (Figures 2C and 2D), confirming that AAR activation by simvastatin results from inhibition of the mevalonate pathway. In support of the transcriptomic findings, simvastatin induced phosphorylation of GCN2 and eIF2α in SK-N-AS and SMS-KCNR neuroblastoma cells (Figure 2E).

**Figure 2.**
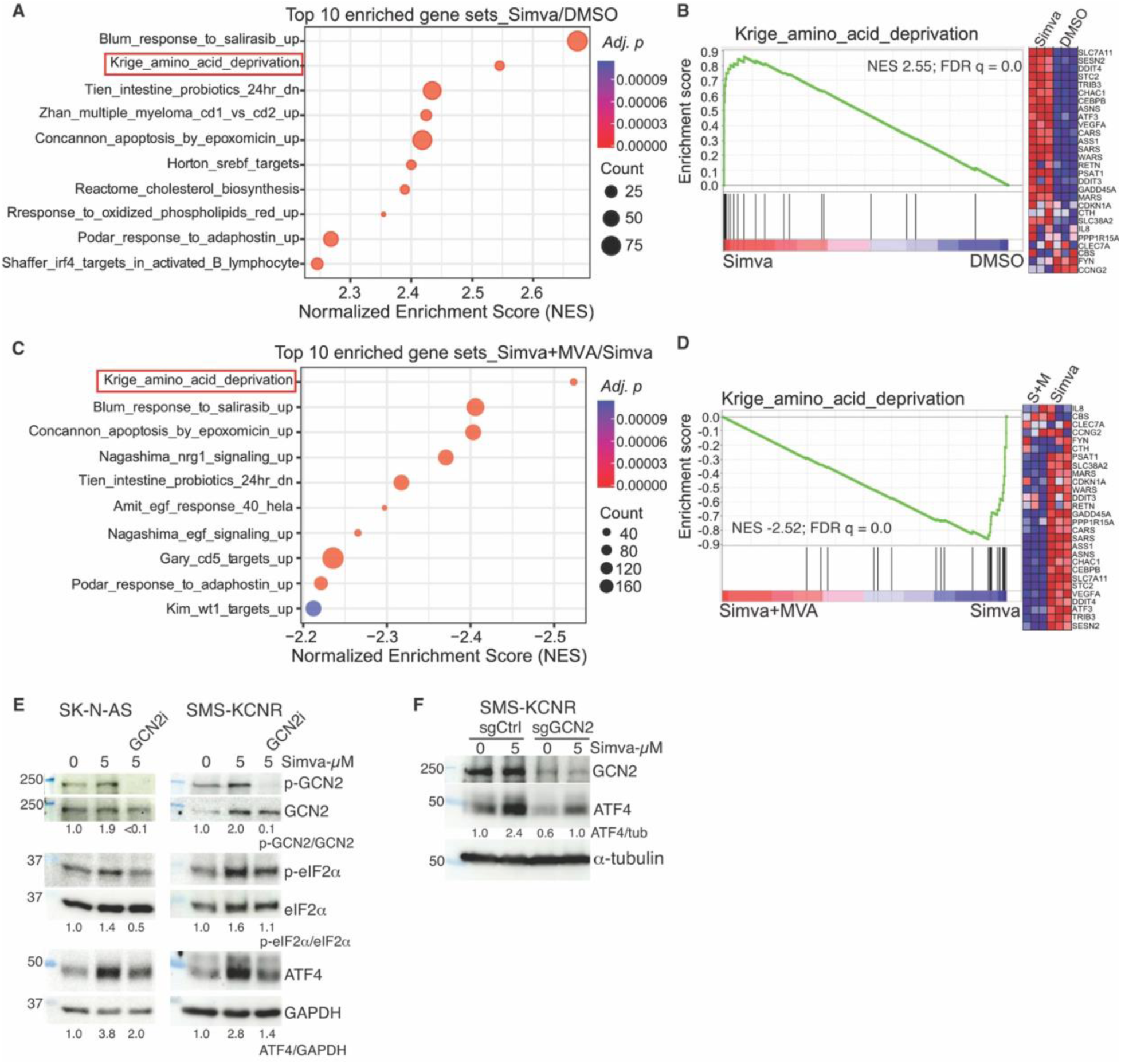
Blocking the mevalonate pathway activates the amino acid deprivation response. (A) Top 10 enriched gene sets in GSEA comparing simvastatin versus DMSO-treated SMS-KCNR cells. (B) GSEA enrichment plot and heatmap showing significant induction of the AAR gene set in SMS-KCNR cells treated with simvastatin compared with DMSO. (C) Top 10 enriched gene sets in GSEA comparing simvastatin + MVA versus simvastatin-treated SMS-KCNR cells. (D) GSEA enrichment plot and heatmap showing significant suppression of the AAR gene set in SMS-KCNR cells treated with simvastatin + MVA compared with simvastatin alone. (E) Immunoblotting showing inhibition of GCN2 activation (phosphorylation) abrogated simvastatin-induced eIF2α phosphorylation and ATF4 upregulation. GCN2 phosphorylation levels were quantified relative to total GCN2, eIF2α phosphorylation levels relative to total eIF2α, and ATF4 levels relative to GAPDH. (F) Immunoblotting showing GCN2 knockout or knockdown by CRISPR abrogated simvastatin induction of ATF4 protein expression. ATF4 levels were quantified relative to α-tubulin.

To determine whether GCN2 mediates simvastatin-induced ATF4 expression, we used a combination of pharmacological and genetic approaches. We first validated the specificity of a GCN2 inhibitor (GCN2i) ^23^ in SK-N-AS cells treated with histidinol (HisOH, an AAR inducer) or tunicamycin (Tm, an ER stress inducer) ^24,25^. GCN2i abrogated ATF4 induction by HisOH but not by Tm, whereas a PERK inhibitor ^26^ blocked ATF4 induction by Tm but not by HisOH (Figure S2), confirming their pathway specificity. Treatment with GCN2i inhibited simvastatin-induced phosphorylation of GCN2 and attenuated ATF4 induction (Figure 2E). Moreover, GCN2 knockout markedly reduced simvastatin-induced ATF4 expression (Figure 2F).

Together, these findings demonstrate that blockade of the mevalonate pathway activates the GCN2-dependent AAR, leading to ATF4 upregulation.

### The mevalonate pathway sustains amino acid biosynthesis

To determine whether amino acid depletion could account for statin-induced AAR, we performed targeted metabolomic profiling of SMS-KCNR neuroblastoma cells treated with DMSO or simvastatin (10 µM, 24 h). Simvastatin treatment significantly decreased the abundance of aspartate, serine, and histidine (Figure 3A and Table S3). Aspartate and serine are non-essential amino acids that can be synthesized by the cell, whereas histidine is an essential amino acid requiring import.

**Figure 3.**
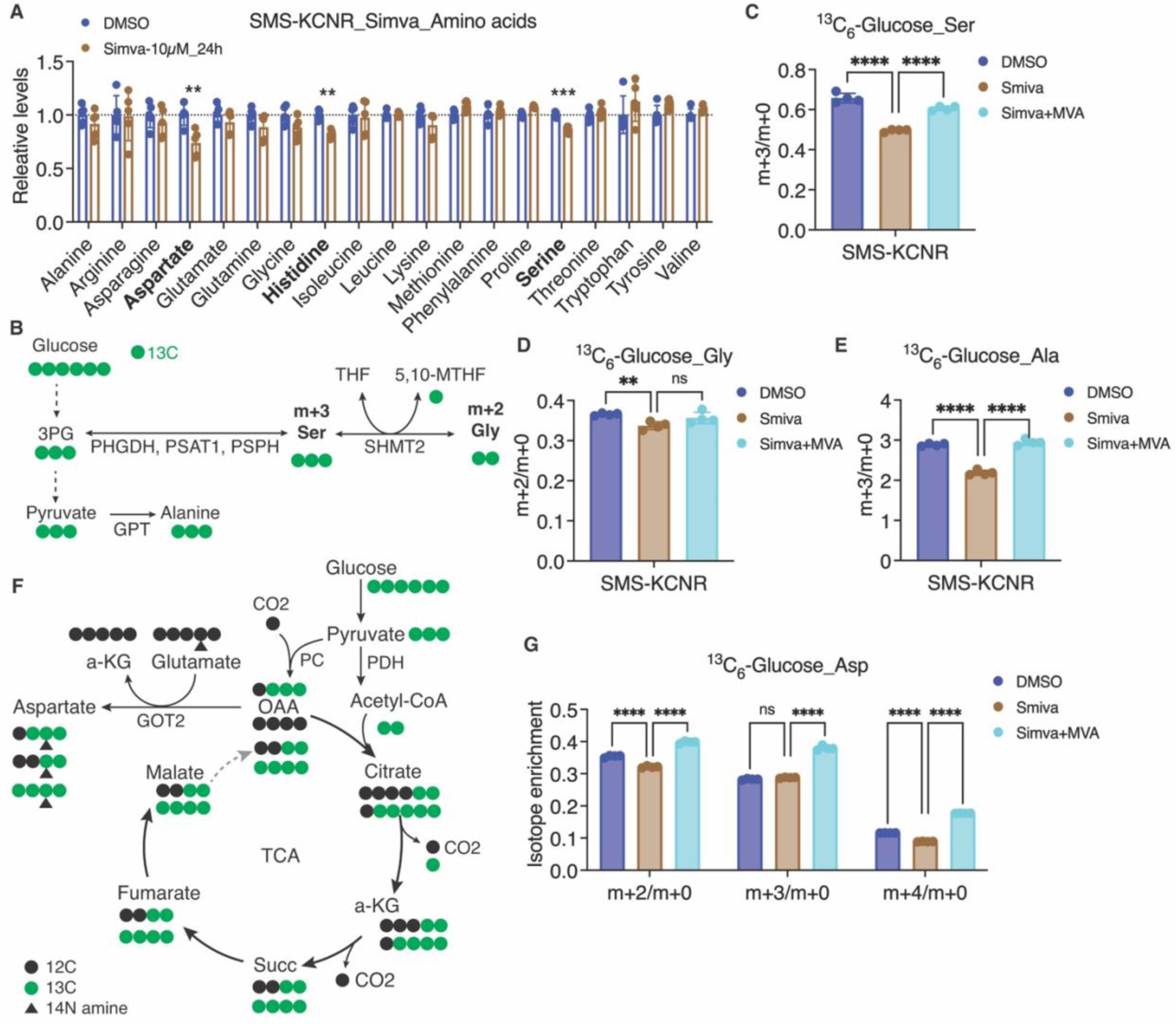
The mevalonate pathway sustains amino acid biosynthesis. (A) Targeted metabolomics analysis of relative intracellular amino acid levels in SMS-KCNR cells treated with DMSO or 10 µM simvastatin for 24 h. (B) Schematic of ^13^C_6_-glucose tracing into serine, glycine and alanine through glycolysis and the serine-glycine synthesis pathway. (C–E) Fractional enrichment of ^13^C-labeled serine (m+3), glycine (m+2), and alanine (m+3) relative to unlabeled species (m+0) in SMS-KCNR cells treated with DMSO, simvastatin, or simvastatin + mevalonate (MVA). (F) Schematic of ^13^C_6_-glucose tracing into aspartate through glycolysis and the TCA cycle. (G) Isotopologue distribution of aspartate (m+2, m+3, m+4) from ^13^C_6_-glucose in SMS-KCNR cells under the indicated treatments. Data in (A, C–E, G) are presented as mean ± SD; statistical significance was determined by one-way ANOVA with Tukey’s multiple comparisons test (**p < 0.01, ***p < 0.001, ****p < 0.0001; ns, not significant).

To investigate whether the observed decreases in serine and aspartate levels result from impaired de novo synthesis, we performed ^13^C_6_-glucose tracing and analyzed isotopologue distributions (Table S4). In the serine–glycine biosynthetic pathway (Figure 3B), simvastatin markedly reduced the fractional enrichment of ^13^C-labeled serine (m+3/m+0) and glycine (m+2/m+0) (Figures 3C and 3D). In the alanine labeling pathway, which reflects pyruvate transamination, simvastatin significantly reduced m+3/m+0 alanine enrichment (Figure 3E). Mevalonate supplementation completely rescued these labeling defects, confirming that they result from inhibition of the mevalonate pathway.

Aspartate is synthesized from oxaloacetate via transamination (Figure 3F). Simvastatin treatment significantly decreased m+2/m+0 and m+4/m+0 aspartate enrichment without affecting m+3/m+0 (Figure 3G), a pattern consistent with reduced glucose carbon flux through the tricarboxylic acid (TCA) cycle into aspartate. Mevalonate supplementation fully restored aspartate labeling to control levels.

Together, these findings demonstrate that the mevalonate pathway supports both glycolytic- and TCA cycle–derived amino acid biosynthesis. Inhibition of this pathway depletes specific amino acids by blocking their synthesis from glucose, providing a direct metabolic basis for statin-induced AAR activation.

### The mevalonate pathway supports de novo purine and pyrimidine biosynthesis

Given that de novo purine and pyrimidine biosynthesis requires amino acid precursors, such as aspartate and glycine whose levels were reduced by simvastatin (Figure 3), we next examined whether inhibition of the mevalonate pathway impairs their incorporation into nucleotides (Table S4). Using ^13^C_6_-glucose tracing, we mapped glucose-derived carbon contributions to nucleotide rings. Ribose labeling from ^13^C_6_-glucose was nearly complete (>95%) and unchanged by simvastatin treatment (Figure S3), indicating that any differences in nucleotide labeling arise from altered incorporation of other precursors.

In purine ring biosynthesis, two carbons are contributed by glycine (m+2) and two carbons by one-carbon unit 10-formyl–THF (m+1-2), in addition to the m+5 from ribose, giving rise to purine isotopologues with m+6 to m+9 labeling (Figure 4A). Simvastatin treatment markedly decreased the fractional enrichment of AMP, ADP, ATP, and GTP in the m+6–9 (or m+6–8 for ADP/ATP/GTP) forms (Figures 4B and 4C), consistent with reduced incorporation of glycine and one-carbon units into the purine ring. Mevalonate supplementation restored labeling to control levels, confirming that this defect is a direct consequence of mevalonate pathway inhibition.

**Figure 4.**
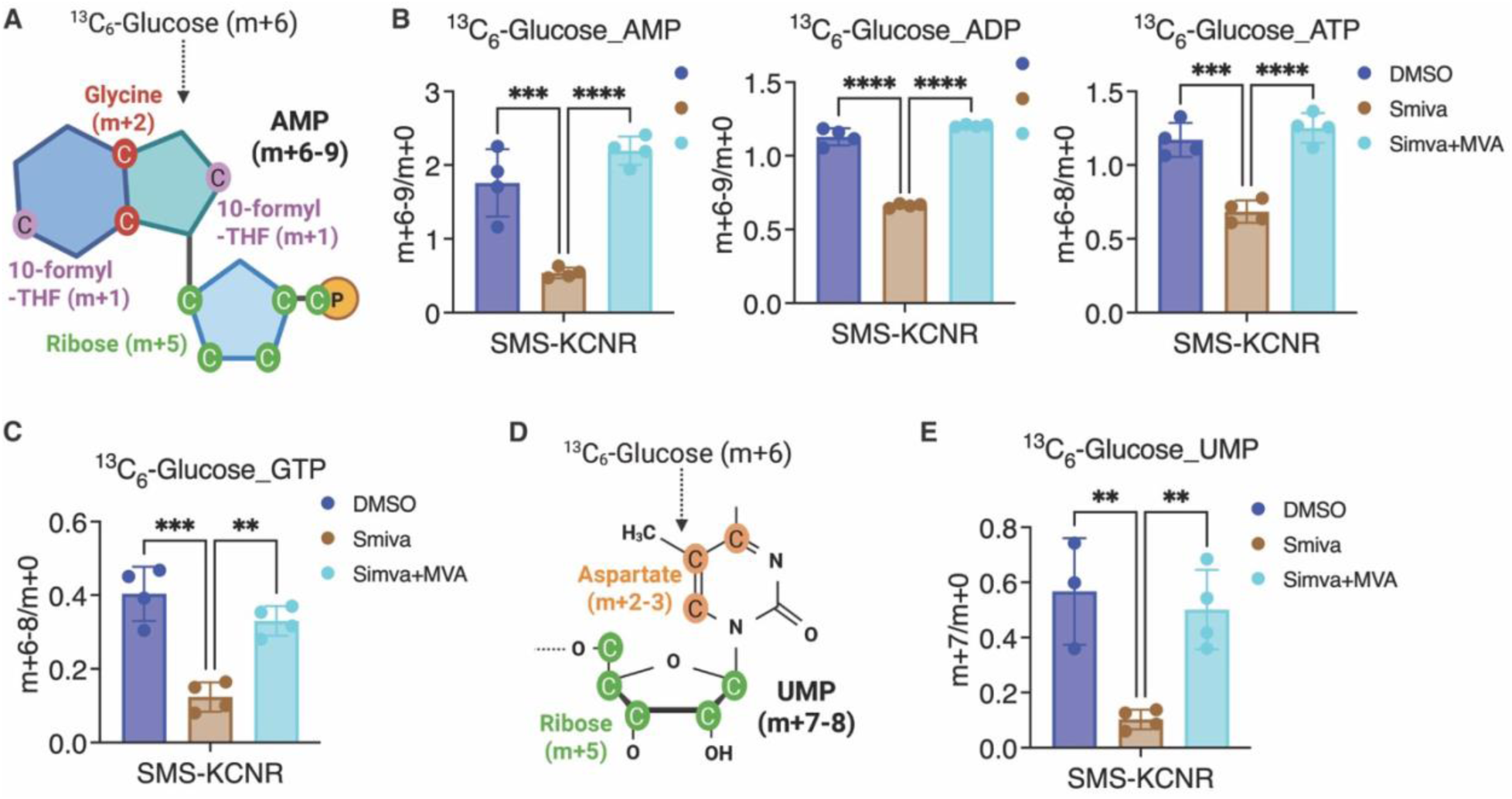
The mevalonate pathway supports de novo purine and pyrimidine biosynthesis. (A) Schematic showing ^13^C_6_-glucose contribution to AMP synthesis through incorporation of glycine-derived carbons (m+2), 10-formyl-THF–derived carbons (m+1), and ribose (m+5). Created with BioRender.com. (B) Fractional enrichment of ^13^C-labeled AMP (m+6–9), ADP (m+6–9), and ATP (m+6–8) relative to unlabeled species (m+0) in cells treated with DMSO, simvastatin, or simvastatin + mevalonate (MVA). Due to co-elution in the LC-MS method, signals labeled as AMP, ADP, and ATP may also include contributions from dGMP, dGDP, dGTP, respectively. However, given the higher abundance and ionization efficiency of ribonucleotides, the detected signals are most likely dominated by AMP, ADP, and ATP. (C) Fractional enrichment of ^13^C-labeled GTP (m+6–8) in the same treatment groups. (D) Schematic showing ^13^C_6_-glucose contribution to UMP synthesis via incorporation of aspartate-derived carbons (m+2–3) and ribose (m+5). Created with BioRender.com. (E) Fractional enrichment of ^13^C-labeled UMP (m+7) relative to m+0 under the indicated treatments. Data in (B, C, E) are presented as mean ± SD; statistical significance was determined by one-way ANOVA with Tukey’s multiple comparisons test (**p < 0.01, ***p < 0.001, ****p < 0.0001).

In the pyrimidine pathway, carbons from aspartate (m+2–3) are incorporated into the pyrimidine ring along with m+5 from ribose, producing UMP isotopologues of m+7–8 (Figure 4D). Simvastatin significantly reduced UMP m+7/m+0 enrichment (Figure 4E), consistent with impaired aspartate incorporation into pyrimidines. Again, mevalonate supplementation rescued the defect.

Collectively, these data indicate that the mevalonate pathway is essential for supplying amino acid precursors and one-carbon units required for de novo purine and pyrimidine biosynthesis.

### The mevalonate pathway sustains NAD⁺ regeneration by the mitochondrial ETC

The mevalonate pathway is required for the synthesis of ubiquinone (coenzyme Q), a lipid-soluble electron carrier that transfers electrons from Complex I and II to Complex III of the mitochondrial electron transport chain (ETC), enabling oxidative phosphorylation and ATP production (Figure 5A) ^5,27^. In addition to driving respiration, ubiquinone maintains redox balance by oxidizing NADH to regenerate NAD⁺, a cofactor required for the de novo biosynthesis of serine and aspartate (Figure 5B) ^28–31^. Statins are known to reduce cellular ubiquinone levels ^32–35^, potentially impairing NAD⁺ regeneration and thereby limiting NAD⁺-dependent biosynthetic processes.

**Figure 5.**
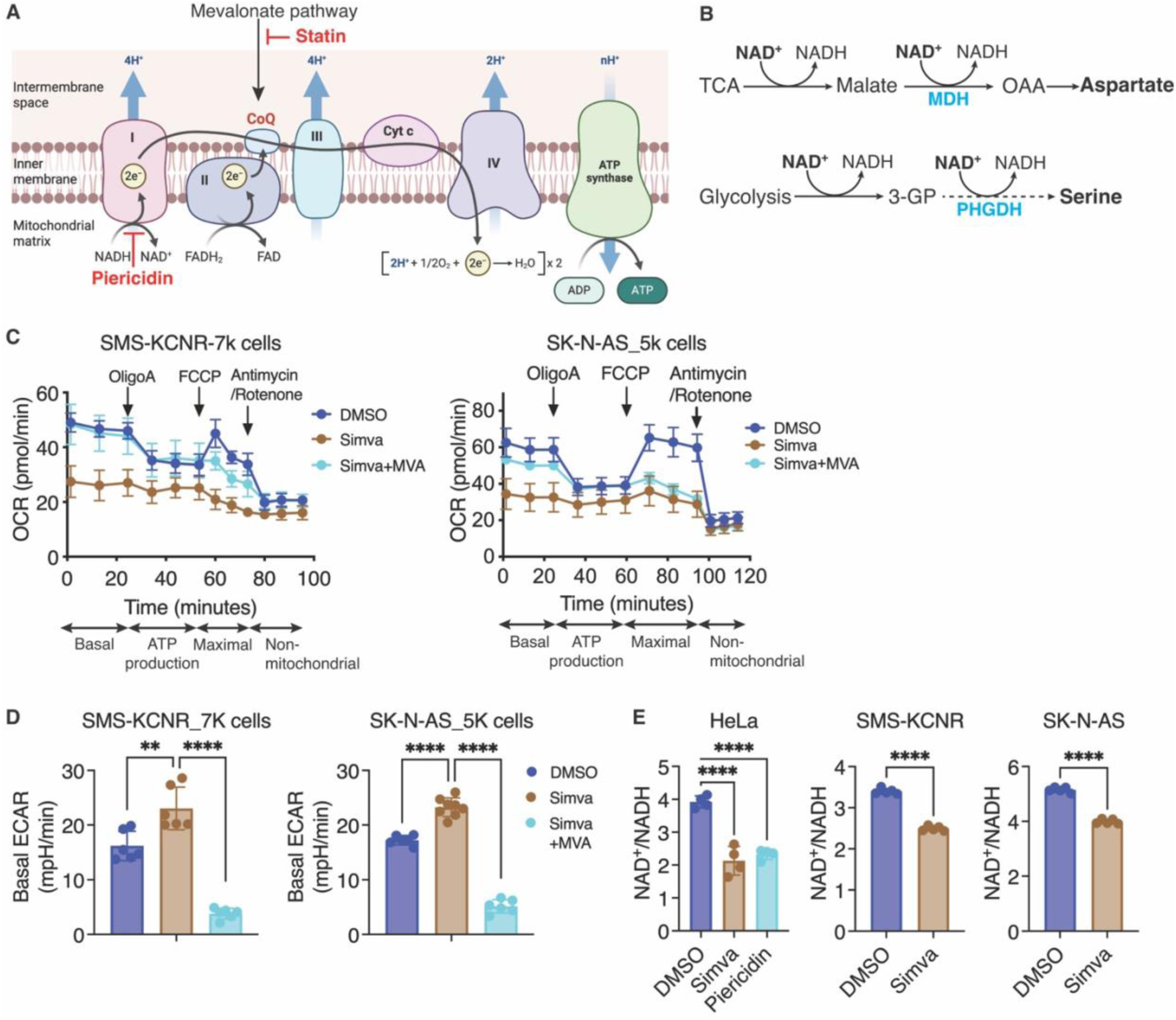
The mevalonate pathway sustains NAD⁺ regeneration by the mitochondrial ETC. (A) Schematic of the mitochondrial ETC showing the mevalonate pathway in CoQ biosynthesis and NAD⁺ regeneration. Statins inhibit HMGCR, reducing CoQ levels and impairing electron transfer from Complex I and II to Complex III. Piericidin inhibits Complex I. Created with BioRender.com. (B) Diagram illustrating NAD⁺-dependent reactions in the TCA cycle for aspartate synthesis and in glycolysis and the serine synthesis pathway for serine production. MDH, malate dehydrogenase. (C) Oxygen consumption rate (OCR) measurements in SMS-KCNR and SK-N-AS cells treated with DMSO, simvastatin (Simva), or simvastatin + mevalonate (Simva+MVA), showing basal respiration, ATP-linked respiration, maximal respiration (FCCP), and non-mitochondrial respiration (antimycin A/rotenone). (D) Basal extracellular acidification rate (ECAR) in SMS-KCNR and SK-N-AS cells under the indicated treatments. (E) Cellular NAD⁺/NADH ratios in HeLa, SMS-KCNR, and SK-N-AS cells after treatment with DMSO, simvastatin, or Piericidin (HeLa only). Data are presented as mean ± SD; statistical significance was determined by one-way ANOVA with Tukey’s multiple comparisons test or unpaired two-tailed t-test (**p < 0.01, ****p < 0.0001).

In support of this model, and consistent with previous reports ^34,35^, simvastatin treatment markedly suppressed basal oxygen consumption rate (OCR), ATP-linked respiration, and maximal respiratory capacity compared with DMSO control (Figure 5C). These defects were largely reversed by mevalonate supplementation, confirming that they result from inhibition of the mevalonate pathway. In parallel, simvastatin treatment increased basal extracellular acidification rate (ECAR) in both SMS-KCNR and SK-N-AS cells (Figure 5D). Elevated basal ECAR reflects increased glycolysis activity, suggesting that simvastatin induces a compensatory shift toward glycolysis.

Direct measurement of the cellular NAD⁺/NADH ratio showed that simvastatin significantly reduced NAD⁺ availability in HeLa, SMS-KCNR, and SK-N-AS cells, with an effect comparable to that of the Complex I inhibitor piericidin in HeLa cells (Figure 5E). These findings indicate that the mevalonate pathway is essential for supporting ETC function and sustaining NAD⁺ regeneration. The resulting redox defect from statin treatment likely contributes to impaired serine and aspartate biosynthesis, both of which require NAD⁺-dependent enzymatic steps (Figure 5B).

### Ubiquinone activity is required for sustaining amino acid pools

To directly test whether ubiquinone supports amino acid biosynthesis, we used complementary approaches to inhibit its production independently of statins. COQ2 is a key enzyme in the ubiquinone biosynthetic pathway that catalyzes the prenylation of 4-hydroxybenzoate (4-HB) with a polyprenyl pyrophosphate side chain. This reaction represents the first committed step in ubiquinone synthesis, linking the aromatic precursor derived from tyrosine to the isoprenoid tail generated by the mevalonate pathway (Figure 6A).

**Figure 6.**
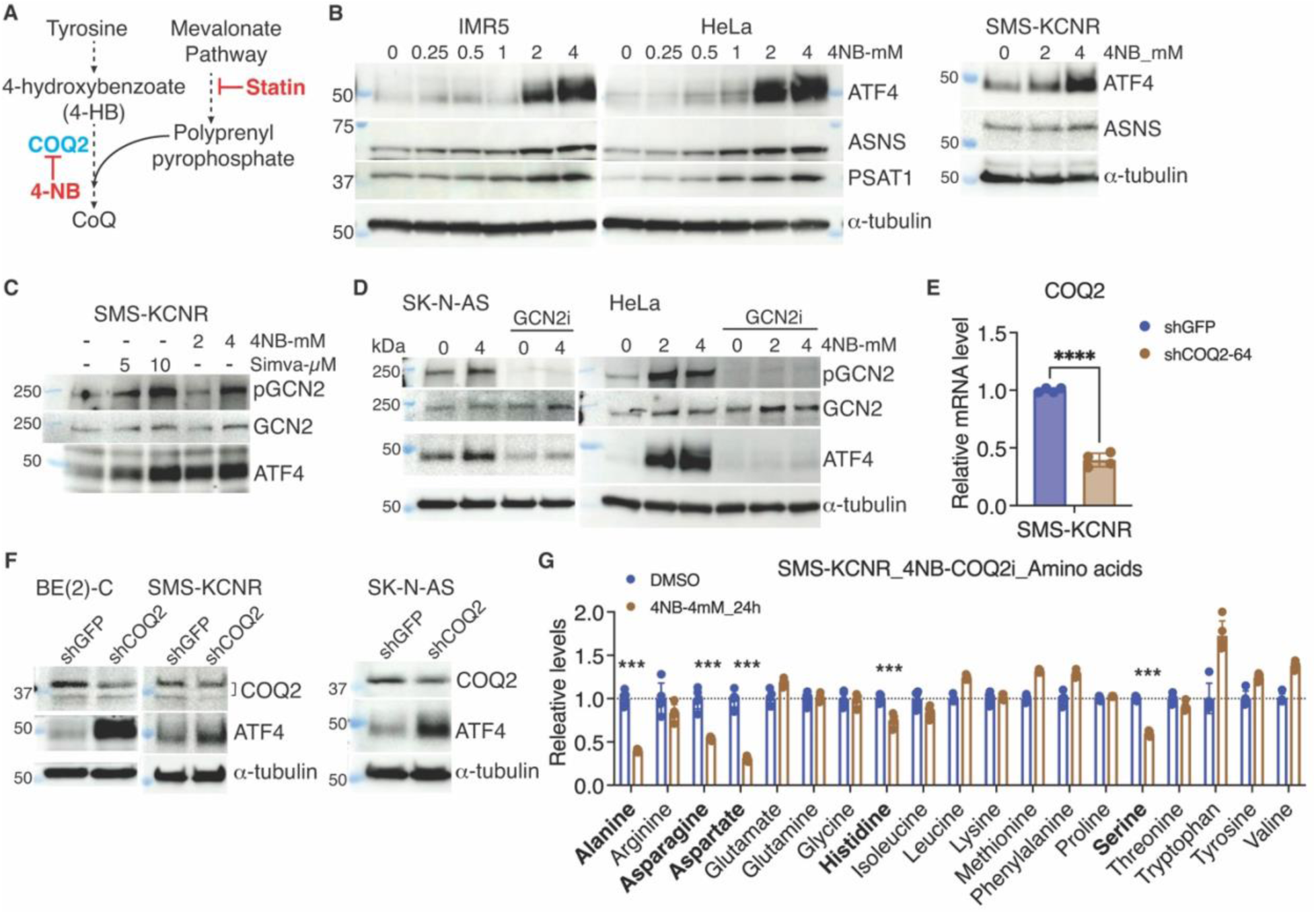
Ubiquinone activity is required for sustaining amino acid pools. (A) Schematic of the ubiquinone (CoQ) biosynthetic pathway. COQ2 catalyzes the condensation of 4-hydroxybenzoate (4-HB) with a polyprenyl pyrophosphate derived from the mevalonate pathway. Statins block the upstream mevalonate pathway, whereas 4-nitrobenzoate (4-NB) competitively inhibits COQ2. (B) Immunoblot analysis of ATF4, ASNS, and PSAT1 in IMR5, HeLa, and SMS-KCNR cells treated with increasing concentrations of 4-NB (0–4 mM) for 24 h. α-tubulin served as a loading control. (C) Immunoblot analysis of p-GCN2, total GCN2, and ATF4 in SMS-KCNR cells treated with indicated concentrations of 4-NB or simvastatin for 24 h. (D) Immunoblot analysis of p-GCN2, GCN2, and ATF4 in SK-N-AS and HeLa cells treated with indicated concentrations of 4-NB for 24 h in the presence or absence of GCN2i. α-tubulin served as a loading control. (E) qRT-PCR analysis of shRNA-mediated knockdown of COQ2 mRNA expression in SMS-KCNR cells (mean ± SD, ****P < 0.0001). (F) Immunoblot analysis of COQ2 and ATF4 in BE(2)-C, SMS-KCNR, and SK-N-AS cells expressing shGFP or shCOQ2. α-tubulin served as a loading control. (G) Targeted metabolomics analysis of relative intracellular amino acid levels in SMS-KCNR cells treated with DMSO or 4NB (4 mM, 24 h). Data represent mean ± SD; ***P < 0.001 by unpaired two-tailed t-test.

To inhibit COQ2, we used 4-nitrobenzoate (4-NB), a structural analog of 4-HB that functions as a competitive inhibitor ^36^. Similar to statin treatment, 4-NB robustly induced ATF4 and its downstream targets ASNS and PSAT1 in neuroblastoma cell lines and HeLa cells (Figures 6B–D and S4), and triggered GCN2 phosphorylation (Figures 6C and 6D). Pharmacological inhibition of GCN2 markedly reduced 4-NB–induced ATF4 expression (Figures 6D and S4), confirming that COQ2 inhibition activates the ATF4-dependent AAR through a mechanism requiring GCN2. Likewise, shRNA-mediated COQ2 knockdown induced ATF4 in multiple cell lines (Figures 6E and 6F), further linking impaired ubiquinone biosynthesis to AAR activation.

Metabolomic analysis revealed that 4-NB treatment significantly lowered steady-state levels of serine, aspartate, and histidine, and also reduced alanine and asparagine levels (Figure 6G and Table S3). Asparagine is synthesized from aspartate and alanine from pyruvate. Thus, the reduction in asparagine and alanine likely reflects diminished availability of their respective precursors, consistent with impaired aspartate and pyruvate metabolism (potentially resulting from a shift toward glycolysis).

These findings establish a specific requirement for ubiquinone in maintaining amino acid pools through its role in sustaining mitochondrial NAD⁺ regeneration.

### Impairing NAD^+^ regeneration is a mechanism for the anti-growth activity of simvastatin

Recent evidence indicates that ubiquinone plays a critical role in sustaining tumor growth ^37^. Because statins block ubiquinone synthesis, we tested whether reduced NAD⁺ regeneration contributes to their anti-proliferative effects. We generated neuroblastoma cell lines expressing *Lactobacillus brevis* NADH oxidase (LbNOX) targeted to either the cytosol or mitochondria (mtNOX) ^38^ (Figure 7A). LbNOX catalyzes NADH oxidation to NAD⁺ using molecular oxygen (Figure 7B), thereby bypassing the mitochondrial ETC. Expression of LbNOX restored the NAD^+^/NADH ratio (Figure 7C) and rescued proliferation (Figure 7D) following simvastatin treatment.

**Figure 7.**
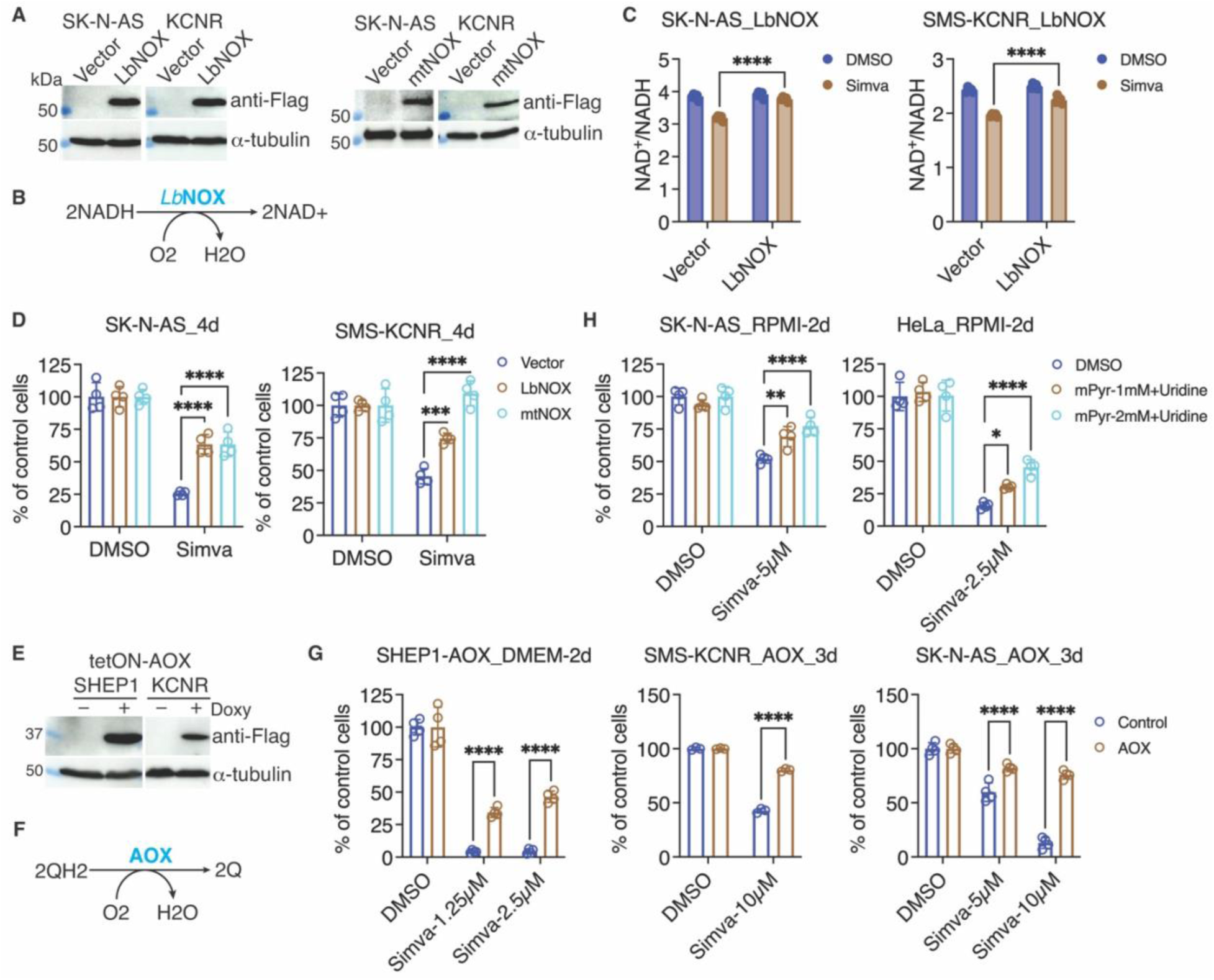
NAD⁺ regeneration or alternative electron transfer restores statin-impaired cell growth. (A) Immunoblot verification of Flag-tagged LbNOX or mitochondrial-targeted LbNOX (mtNOX) expression in SK-N-AS and SMS-KCNR cells. α-tubulin served as a loading control. (B) Schematic of LbNOX-catalyzed oxidation of NADH to NAD⁺ using molecular oxygen as an electron acceptor. (C) Intracellular NAD⁺/NADH ratio in SK-N-AS and SMS-KCNR cells expressing LbNOX following treatment with DMSO or simvastatin. (D) Cell viability of SK-N-AS and SMS-KCNR cells (4 days) expressing vector, LbNOX, or mtNOX and treated with DMSO or simvastatin. (E) Immunoblot confirmation of doxycycline (Doxy)-inducible, Flag-tagged AOX expression in SHEP1 and SMS-KCNR cells. (F) Schematic of AOX-mediated oxidation of ubiquinol (QH₂) to ubiquinone (Q) with oxygen as the electron acceptor. (G) Cell viability of SHEP1, SMS-KCNR, and SK-N-AS cells without (control) or with Doxy-inducible expression of AOX following simvastatin treatment. (H) Cell viability of SK-N-AS and HeLa cells cultured in RPMI medium and treated with simvastatin in the presence of 1 mM or 2 mM methyl-pyruvate (mPyr) plus 0.1 µM uridine. Data are presented as mean ± SD; *p < 0.05, **p < 0.01, ***p < 0.001, ****p < 0.0001 by two-way ANOVA with multiple comparisons.

We further tested this model using an alternative ETC bypass system by introducing the alternative oxidase (AOX) from *Ciona intestinalis* into neuroblastoma cells (Figure 7E). AOX transfers electrons directly from ubiquinol to molecular oxygen (Figure 7F) ^37,39^, thereby circumventing ETC blockades. AOX expression similarly mitigated simvastatin-induced growth inhibition (Figure 7G).

Previous studies have shown that combined pyruvate and uridine supplementation rescues cells lacking a functional ETC ^40^. Pyruvate supports TCA cycle–derived aspartate synthesis ^28,29^ and promotes NAD^+^ regeneration ^41,42^, while uridine can bypass the requirement for de novo pyrimidine biosynthesis, which depends on ETC-driven DHODH activity ^43^. Consistent with our model, supplementation with both methyl-pyruvate and uridine significantly rescued the growth inhibitory effect of simvastatin (Figure 7H).

Collectively, these findings support a model in which inhibition of electron flow through the ETC, and the resulting disruption of NAD⁺ regeneration, is an important mechanism contributing to the anti-proliferative activity of statins.

## Discussion

Our study identifies the mevalonate pathway as a key regulator of amino acid biosynthesis through its role in sustaining mitochondrial NAD⁺ regeneration. Inhibition of this pathway by statins impairs oxidative phosphorylation, lowers the cellular NAD⁺/NADH ratio, and reduces the synthesis of serine and aspartate, leading to activation of the GCN2–eIF2α–ATF4 axis that drives the AAR. The resulting depletion of the serine and aspartate pools disrupts purine and pyrimidine nucleotide synthesis. These effects result from diminished production of ubiquinone, a mevalonate-derived electron carrier essential for mitochondrial respiration. Restoring NAD⁺ levels through expression of NADH-oxidizing enzymes attenuates the growth-inhibitory effects of statins, establishing a mechanistic link between mevalonate pathway activity, redox balance, and amino acid homeostasis. Collectively, these findings uncover a metabolic role for the mevalonate pathway in coupling mitochondrial electron transport to amino acid biosynthesis and identify redox imbalance as a key driver of the anti-proliferative effects of statins in cancer cells.

Amino acid biosynthesis is tightly integrated with cellular energy metabolism and central carbon flux, requiring coordinated input from glycolysis, the TCA cycle, and mitochondrial electron transport ^44–46^. Among the cofactors bridging these metabolic processes, NAD⁺ plays a central role in maintaining redox balance and driving dehydrogenase-catalyzed reactions that support amino acid production ^47–49^. Recent studies have underscored the importance of NAD⁺ regeneration in sustaining anabolic pathways, particularly in proliferating cells where high demand for aspartate and serine creats a strong reliance on mitochondrial respiration. Inhibition of the ETC depletes the cellular NAD⁺ pool and restricts aspartate synthesis, thereby suppressing cell growth ^28,29,31,50^. Likewise, a reduced NAD^+^/NADH ratio impairs serine biosynthesis and suppresses cell proliferation ^30,51^. These findings collectively highlight NAD⁺ availability as a critical determinant of amino acid biosynthetic capacity.

Our findings expand this framework by positioning the mevalonate pathway as a critical upstream regulator of NAD⁺ regeneration, acting through its role in ubiquinone biosynthesis. Ubiquinone is an essential electron carrier in the ETC, shuttling electrons from Complex I and II to Complex III, thereby driving oxidative phosphorylation and enabling NADH oxidation to regenerate NAD^+ 5,27^. We show that statin-mediated inhibition of the mevalonate pathway reduces ubiquinone levels, resulting in suppression of de novo serine and aspartate synthesis. This in turn limits serine-derived glycine and one-carbon units, as well as aspartate, which are essential precursors for purine and pyrimidine biosynthesis. Traditionally, the mevalonate pathway has been studied in the context of cholesterol synthesis and protein prenylation ^1,2,52–54^. Our findings broaden this view by placing the mevalonate pathway at the intersection of lipid metabolism and redox-dependent amino acid and nucleotide biosynthesis.

A key implication is that impaired NAD⁺ regeneration is not merely a secondary consequence of mevalonate pathway inhibition but constitutes a direct mechanism contributing to statin-induced growth suppression in cancer cells. We show that ectopic expression of LbNOX, a bacterial NADH oxidase that oxidizes NADH to NAD⁺ using molecular oxygen, restored the NAD⁺/NADH ratio and rescue proliferation in simvastatin-treated neuroblastoma cells. Similarly, expression of AOX, which bypasses ubiquinone and Complex III by transferring electrons directly from ubiquinol to oxygen, mitigated the growth defect. These rescue experiments provide strong genetic evidence that ETC dysfunction and consequent NAD⁺ deficiency are causally linked to anti-proliferative activity of statins.

In summary, our work establishes a pivotal role for the mevalonate pathway in maintaining mitochondrial function, redox balance, and amino acid production. By defining how statin-induced disruption of NAD⁺ regeneration leads to amino acid insufficiency and growth inhibition, we provide a mechanistic framework in which lipid metabolism supports mitochondrial energy and biosynthetic functions that drive de novo amino acid and nucleotide synthesis. These insights not only expand the biological scope of the mevalonate pathway but also suggest therapeutic opportunities, either by targeting this pathway in tumors with high amino acid demand or by exploiting its dependence on ubiquinone-mediated NAD⁺ regeneration to sensitize cancer cells to metabolic stress.

## ACKNOWLEDGEMENTS

We thank Wilson Landon and Dr. Stephen Barnes at the UAB Targeted Proteomics & Metabolomics Lab for amino acid analysis, and Dr. Peng Gao for assistance with stable isotope metabolomics conducted at the Metabolomics Core Facility of the Robert H. Lurie Comprehensive Cancer Center, Northwestern University (supported by NCI CCSG P30 CA060553). S.S. is supported by NIH R01CA200653, DoD HT94252410558, and Department of Veterans Affairs I01BX002930. This work was supported by NIH R01CA190429 and R01CA236890 to H.-F.D.

## AUTHOR CONTRIBUTIONS

T.T.N., and H-F.D. conceived and designed the study with contributions from M.B., J.D., and S.S. T.T.N., M.B., J.D., and H.-F.D. performed the experiments, T.T.N., M.B., J.D., and H-F.D. analyzed data with contributions from S.S. T.T.N. and H-F.D. wrote the paper with contributions from J.D., M.B. H-F.D. supervised and provided funding for the project. All authors read the manuscript and approved its contents.

## STAR*METHODS

### KEY RESOURCES TABLE

| REAGENT or RESOURCE | SOURCE | IDENTIFIER |
| --- | --- | --- |
| **Antibodies** |  |  |
| Mouse anti-α-tubulin (clone B-5-1-2) | Sigma-Aldrich | T5168 |
| Mouse anti-β-actin | Invitrogen | MA5-15739 |
| Rabbit anti-ACSS2 | Cell Signaling | #3658 |
| Rabbit anti-ASNS | Proteintech | 14681-1-AP |
| Rabbit anti-ATF3 | Cell Signaling | #33593 |
| Rabbit anti-ATF4 | Cell Signaling | #11815 |
| Rabbit anti-COQ2 | Prointech | 30909-1-AP |
| Mouse anti-eIF2α (D-3) | Santi Cruz | sc-133132 |
| Rabbit anti-phospho-eIF2α (Ser51) | Cell Signaling | #9721 |
| Rabbit anti-GAPDH (F-335) | Santa Cruz | sc-25778 |
| Mouse anti-GCN2 (F-7) | Santa Cruz | Sc-374609 |
| Rabbit anti-phospho-GCN2 (T899) | Abcam | ab75836 |
| Rabbit anti-PSAT1 | Novus | 21020002 |
| Rabbit anti-SLC1A5/ASCT2 (V501) | Cell Signaling | #5345 |
| Rabbit anti-SLC7A5 | Cell Signaling | #5347 |
| Rabbit anti-SLC7A11/xCT (D2M7A) | Cell Signaling | #12691 |
| Goat anti-mouse IgG HRP | Jackson ImmunoResearch | 115-035-146 |
| Goat anti-rabbit IgG HRP | Jackson ImmunoResearch | 111-035-046 |
| **Chemicals and Inhibitors** |  |  |
| <sup>13</sup>C<sub>6</sub>-glucose | Sigma-Aldrich | 389374-CONF |
| Cholesterol-water soluble | Sigma-Aldrich | C4951 |
| DMSO | Fisher Scientific | BP231 |
| Farnesyl Pyrophosphate (FPP) | Cayman | 63250 |
| GCN2iB | MedChemExpress | HY-112654 |
| GeranylGeranyl Pyrophosphate (GGPP) | Cayman | 63330 |
| Histidinol | Sigma-Aldrich | H6647 |
| Mevalonolactone (Mevalonic acid lactone) | Sigma-Aldrich | M4667 |
| PERKi, GSK2606414 | Sigma-Aldrich | 516535 |
| Pitavastatin | Selleckchem | S1759 |
| Simvastatin | Selleckchem | S1792 |
| Trizol | Thermo Fisher Scientific | 15596026 |
| Tunicamycin | Sigma-Aldrich | T7765 |
| **Critical Commercial Assays** |  |  |
| 2X SYBR green qPCR master mix | Bimake | B21203 |
| iScript Advanced cDNA Synthesis Kit for RT-qPCR | Bio-Rad | 172-5038 |
| Bio-Rad Protein Assay Kit II | Bio-Rad | 5000002 |
| Clarity Western ECL Substrate Kit | Bio-Rad | 1705061 |
| NAD<sup>+</sup>/NADH-Glo Assay Kit | Promega | G9071 |
| **Deposited data** |  |  |
| Microarray data | 13 | GEO GSE215949 |
| **Experimental Models: Cell Lines** |  |  |
| 293FT | Thermo Fisher | R70007 |
| 293T (ATF4 knockout) | 55 | N/A |
| 293T Lenti-X | TaKaRa | 632180 |
| BE(2)-C | ATCC | CRL-2268 |
| HCT116 | ATCC | CCL-247 |
| HeLa | ATCC | CCL-2 |
| IMR5 | 56 | N/A |
| SHEP1 | 57 | N/A |
| SK-N-AS | ATCC | CRL-2137 |
| SMS-KCNR | Children's Oncology Group<br>https://www.cccells.org/cellreqs-all.php | N/A |
| **Oligonucleotides** |  |  |
| qRT-PCR primers | This paper | See Table S5 |
| **DNA constructs** |  |  |
| pCDH-CMV-MCS-EF1-puro | SBI System Biosciences | CD510B-1 |
| pCW57.1 AOX-FLAG | Addgene | Plasmid #177984 |
| pUC57-LbNOX | Addgene | Plasmid #75285 |
| pUC57-mitoLbNOX | Addgene | Plasmid #74448 |
| pLKO.1-shGFP | Addgene | Plasmid #12273 |
| pLKO.1-shCOQ2-64 | Millipore Sigma | TRCN0000072564 |
| pLKO.1-shHMGCR-14 | Millipore Sigma | TRCN0000233114 |
| pLKO.1-shHMGCR-15 | Millipore Sigma | TRCN0000233115 |
| pLP1 | Thermo Fisher | K497500 |
| pLP2 | Thermo Fisher | K497500 |
| pLP/VSVG | Thermo Fisher | K497500 |
| **Software and Algorithms** |  |  |
| Prism for Mac OS X version 10.5.0 | GraphPad | http://www.graphpad.com/ |
| ImageJ2 (version 2.16.0/1.54p) | NIH | https://imagej.net/ij |
| DAVID Bioinformatics Resources 6.8 | NIAID/NIH | https://david.ncifcrf.gov/home.jsp |
| GSEA | Broad Institute | https://www.gsea-msigdb.org/gsea/index.jsp |
| Partek Genomics Suite | Partek | https://www.partek.com/ |
| Other |  |  |

### Resource availability Lead contact

Further information and requests for resources and reagents should be directed to and will be fulfilled by the Lead Contact, Dr. Han-Fei Ding.

### Materials availability

This study did not generate new unique reagents.

## EXPERIMENTAL MODEL AND SUBJECT DETAILS

### Cell lines

All neuroblastoma cell lines and their culture conditions have been described in detail previously ^58^. 293FT (Thermo Fisher R70007), 293T-ATF4 KO ^55^, 293T Lenti-X (TaKaRa 632180), HeLa (ATCC CCL-2), and HCT116 (ATCC CCL-247) were cultured in DMEM (HyClone SH30022) supplemented with 10% fetal bovine serum (FBS; Atlanta Biologicals S11050). All cell lines underwent authentication via short tandem repeat (STR) profiling (ATCC). Following authentication, large batches of frozen stocks were prepared to minimize cross-contamination risk. Cells were used within 10 passages after thawing and confirmed mycoplasma-free by DAPI staining every three months. Neuroblastoma cell lines were routinely validated by immunoblotting and immunofluorescence for high-level nuclear expression of MYCN and the specific neuroblastoma marker PHOX2B ^59^. Cell growth and viability were determined by trypan blue exclusion assay or MTT assay.

## METHOD DETAILS

### Drugs and small molecules

Simvastatin (Selleckchem, S1792), pitavastatin (Selleckchem S1759), GCN2iB (MedChemExpress, HY-112654), and GSK2606414 PERK inhibitor (Millipore Sigma, 1337531-89-1) were dissolved in DMSO (Thermo Fisher Scientific, BP231), aliquoted, and stored at −80 °C. FPP (Cayman 63250) and GGPP (Cayman 63330) were supplied as a solution of methanol:NH_4_OH (70:30) and stored at -20°C. Mevalonolactone (Sigma-Aldrich M4667) was diluted in H_2_O and stored at -20°C. Cells were treated with individual agents or combinations the indicated concentrations and durations before collection for viability assays (trypan blue or MTT), qRT-PCR, or immunoblotting. For AAR or ER stress induction, cells were treated with vehicle controls (H2O for HisOH and DMSO for tunicamycin), 5 mM Histidinol (HisOH, Sigma-Aldrich H6647), or 2 µg/ml tunicamycin (Sigma-Aldrich T7765) for 8 h before immunoblotting.

### Overexpression and RNA interference

Lentiviral CRISPR/Cas9 sgRNA constructs targeting ATF4 and GCN2 were acquired from Applied Biological Materials. The plasmids pCW57.1_AOX-Flag, pUC57-LbNOX and pUC57-mitoLbNOX were acquired from Addgene. Coding sequences for LbNOX and mitoLbNOX, including a Flag tag, were subcloned into the pCDH lentiviral vector. Lentiviruses were generated in 293T Lenti-X cells using packaging plasmids pLP1 (RRID: Addgene_22614), pLP2, and pLP/VSVG (Thermo Fisher Scientific, K497500). Lentiviral infections were performed following standard protocols.

### Quantitative real-time PCR (qRT-PCR)

Total RNA was extracted using TRIzol reagent (Thermo Fisher Scientific, 15596026). Reverse transcription was performed with the iScript Advanced cDNA Synthesis Kit (Bio-Rad, 172-5038). qRT-PCR was carried out with 2× SYBR Green qPCR Master Mix (Bimake, B21203) on an iQ5 real-time PCR system (Bio-Rad) using gene-specific primers listed in Table S5. Expression levels were normalized to β₂-microglobulin (B2M) mRNA.

### Immunoblotting

Cells were lysed in standard SDS sample buffer, and protein concentration was determined using the Bio-Rad Protein Assay Kit II. Equal amounts (30–50 µg) were separated by SDS-PAGE, transferred to nitrocellulose membranes, and probed with primary antibodies listed in the **Key Resources Table**. HRP-conjugated secondary antibodies included goat anti-mouse IgG (Jackson ImmunoResearch, 115-035-146, RRID:AB_2307392) and goat anti-rabbit IgG (Jackson ImmunoResearch, 111-035-046, RRID:AB_2337939). Signals were detected using Clarity Western ECL (Bio-Rad, 1705061) and imaged with the Amersham ImageQuant 800 (Cytiva) or quantified using ImageJ2 (v2.16.0/1.54p).

### NAD+/NADH assay

NAD⁺/NADH ratios were measured as described previously (6). Briefly, 4×10⁵ cells were seeded in 6-cm dishes and incubated overnight before 6 h drug treatment. Cells were washed with ice-cold PBS, lysed in PBS containing 1% dodecyltrimethylammonium bromide in 0.2 M NaOH, and divided into two aliquots—one untreated and one treated with 1 M ascorbic acid followed by 0.4 M HCl. Samples were heated at 60 °C for 17 min to degrade reduced or oxidized nucleotides selectively. NAD⁺ and NADH levels were quantified using the Promega NAD⁺/NADH-Glo Assay Kit (G9071).

### Targeted metabolomics

SMS-KCNR cells were treated with DMSO, simvastatin (10 µM), or 4-NB (4 mM) for 24 h. After two PBS washes, cells were extracted in methanol at −20°C for 30 min, scraped, and centrifuged at 13,000 rpm for 10 min at 4°C. Supernatants were dried under N₂ gas and reconstituted in 50% methanol. Multiple dilutions ensured measurements were within the standard curve range. Amino acids were quantified using LC-MS/MS as described previously ^60^, and data were analyzed using MultiQuant 3.0.3 (SCIEX).

### Stable isotope flux analysis

SMS-KCNR cells were treated with DMSO, simvastatin (10 µM), or simvastatin (10 µM) plus mevalonate (2 mM) for 24 h, then switched to glucose-free DMEM (Invitrogen, 11966-025) supplemented with 10% dialyzed FBS and 25 mM [U-¹³C₆]-glucose for 10 h. Four biological replicates (∼5×10⁶ cells/sample) per treatment were analyzed.

Metabolites were extracted with 80% methanol at −80°C, dried in a SpeedVac, and stored at −80°C. Pellets were reconstituted in 60% acetonitrile, vortexed, centrifuged, and supernatants analyzed by HPLC-MS/MS. Data acquisition and analysis were performed using Xcalibur 4.1 and TraceFinder 4.1 (Thermo Fisher Scientific) as described previously ^61^.

### Statistics

Data are presented as mean ± SD. Statistical significance was assessed by unpaired two-tailed Student’s *t*-test or one-/two-way ANOVA, as appropriate, using GraphPad Prism 10.5.0 (Mac).

