## Supplemental Data for "The mevalonate pathway couples lipid metabolism to amino acid synthesis via ubiquinone-dependent redox control"

### The mevalonate pathway sustains amino acid synthesis via ubiquinone–driven redox homeostasis

##### Inventory of Supplemental Information

##### Supplemental Figures S1-S7

Figure S1 related to Figure 1
Figure S2 related to Figure 2
Figure S3 related to Figure 4
Figure S4 related to Figure 6

##### Supplemental Tables S1-S6

Table S1 related to Figure 1
Table S2 related to Figure 1
Table S3 related to Figure 3, 6
Table S4 related to Figures 3, 4
Table S5 related to Figures 1, 6

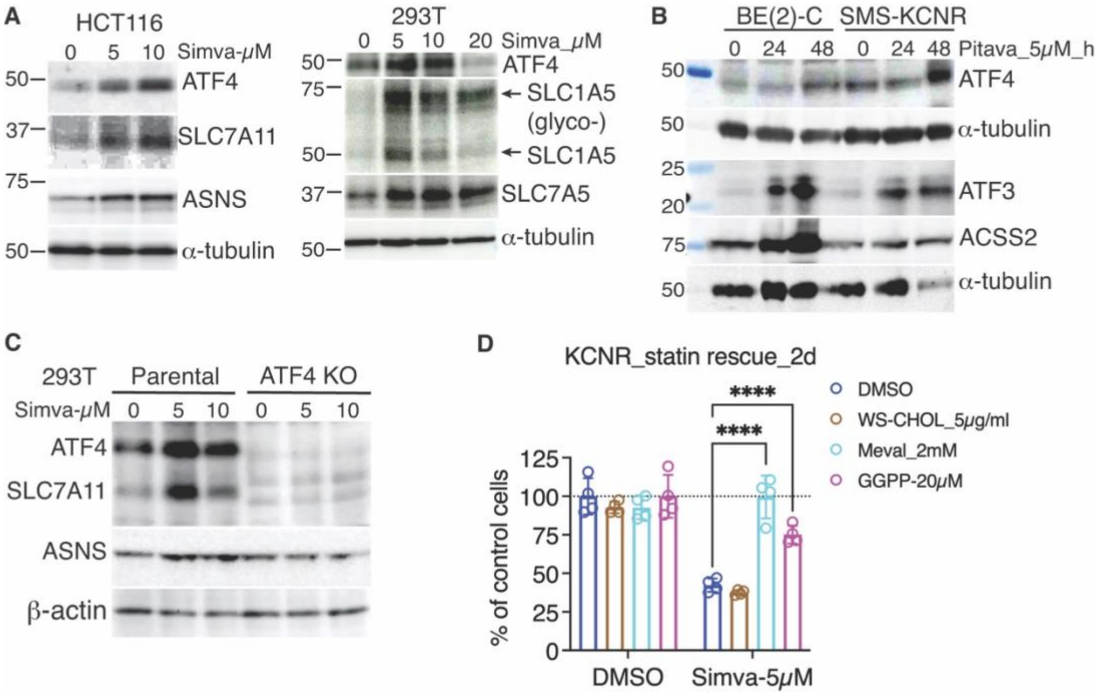


**Figure S1. Statins activate the ATF4-mediated amino acid response pathway.**
(A) Immunoblot analysis of ATF4 and its downstream targets SLC7A11 and ASNS in HCT116 and 293T cells treated with increasing concentrations of simvastatin for 24 h. Glycosylated and non-glycosylated forms of SLC1A5 are indicated.
(B) BE(2)-C and SMS-KCNR neuroblastoma cells were treated with 5 μM pitavastatin for 0, 24, or 48 h. Immunoblotting was performed for ATF4, ATF3, and ACSS2. α-Tubulin served as a loading control.
(C) Immunoblot analysis of ATF4, SLC7A11, and ASNS in parental and ATF4 knockout (KO) 293T cells treated with simvastatin (0, 5, or 10 μM) for 24 h.
(D) Cell viability of SMS-KCNR cells treated with DMSO or 5 μM simvastatin for 48 h, with or without co-treatment with water-soluble cholesterol (5 μg/ml), mevalonate (2 mM), or geranylgeranyl pyrophosphate (GGPP, 20 μM). Data are shown as mean ± SD, n = 4 replicates. ****$p < 0.0001$ by one-way ANOVA with Tukey's test.

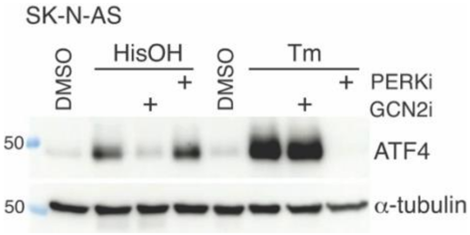


**Figure S2. GCN2 is required for ATF4 induction by amino acid deprivation.** Immunoblot analysis of ATF4 in SK-N-AS cells treated with histidinol (HisOH, 2 mM) or tunicamycin (Tm, 2 μg/ml) for 8 h in the presence or absence of GCN2 inhibitor (GCN2i, 1 μM) or PERK inhibitor (PERKi, 1 μM). α-Tubulin was used as a loading control.

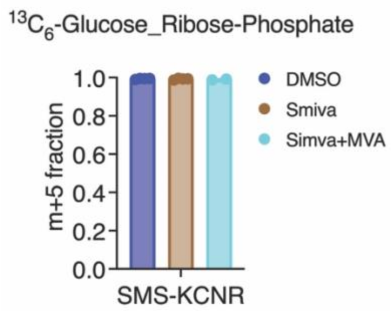


**Figure S3. Statin treatment does not affect ribose-5-phosphate labeling from glucose.**
Fractional enrichment of m+5 ribose-phosphate derived from [U-$^{13}C_6$]-glucose in SMS-KCNR cells treated with DMSO, simvastatin (Simva, 5 μM), or simvastatin plus mevalonate (MVA, 2 mM) for 24 h. Data represent mean ± SD (n = 4 for DMSO and Simva groups; n = 2 for Simva + MVA group).

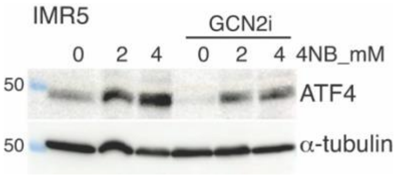


**Figure S4. 4-Nitrobenzoate (4-NB) induces ATF4 in a GCN2-dependent manner.** Immunoblot analysis of ATF4 expression in IMR5 neuroblastoma cells treated with 4-NB (2 or 4 mM) for 24 h, with or without the GCN2 inhibitor (GCN2i, 1μM). α-Tubulin serves as a loading control.
